# Modeling the time-resolved modulations of cardiac activity in rats: A study on pharmacological autonomic stimulation

**DOI:** 10.1101/2024.03.27.586625

**Authors:** Diego Candia-Rivera, Sofia Carrion-Falgarona, Fabrizio de Vico Fallani, Mario Chavez

**Affiliations:** Sorbonne Université, Paris Brain Institute (ICM), CNRS UMR7225, INRIA Paris, INSERM U1127, Hôpital de la Pitié Salpêtrière AP-HP, 75013, Paris, France

**Keywords:** Autonomic nervous system, heart rate variability, dobutamine, Poincare plot

## Abstract

Assessing cardiac dynamics over time is essential for understanding cardiovascular health and its parallel patterns of activity with the brain. We present a methodology to estimate the time-resolved sympathetic and parasympathetic modulations of cardiac dynamics, specifically tailored for the rat heart. To evaluate the performance of our method, we study a dataset comprising spontaneously hypertensive rats and Wistar-Kyoto rats. These rats were administered dobutamine to elicit autonomic dynamics. The results obtained from our method demonstrated accurate time-resolved depiction of sympathetic reactivity induced by dobutamine administration. These responses closely resembled the expected autonomic alterations observed during physical exercise conditions, albeit emulated pharmacologically. We further compared our method with standard measures of low-frequency (LF) and high-frequency (HF) components, which are commonly used, although debated, for sympathetic and parasympathetic activity estimation. The comparisons with LF and HF measures further confirmed the effectiveness of our method in better capturing autonomic changes in rat cardiac dynamics.

Our findings highlight the potential of our adapted method for time-resolved analysis in future clinical and translational studies involving rodents’ models. The validation of our approach in animal models opens new avenues for investigating the relationship between ongoing changes in cardiac activity and parallel changes in brain dynamics. Such investigations are crucial for advancing our understanding of the brain-heart connection, particularly in cases involving neurodegeneration, brain injuries, and cardiovascular conditions.

**Key points:**

– We developed a method for time-resolved estimation of sympathetic and parasympathetic modulations in rat cardiac dynamics, validated against standard LF and HF measures.
– We used a cohort of spontaneously hypertensive rats and Wistar-Kyoto rats, with dobutamine administration to induce autonomic responses.
– Our method accurately depicted time-resolved sympathetic reactivity similar to autonomic changes during physical exercise.
– Our findings suggest potential for future clinical and translational studies on the brain-heart connection, particularly in cardiovascular conditions.

## Introduction

Cardiac activity is regulated in the short term through two synergistic pathways: the sympathetic system, which acts as a positive ionotropic, and the parasympathetic system, with negative ionotropic effect (Ruffolo, 1987; Spyer, 1990; Mahoney *et al*., 2016), being these disruptions typically manifested in the heart rate and rhythmicity (or heart rate variability; HRV) (Macpherson *et al*., 2009). Standard HRV analysis in rats typically relies on 2-3 minute time-averaged measures, such as mean heart rate or standard deviation of R-R intervals (Kuwahara *et al*., 1994; Rowan *et al*., 2007). However, this approach overlooks the rich oscillatory nature of cardiac dynamics, which could provide further insights about the sympathetic and parasympathetic modulations of cardiac dynamics.

Recent advancements in humans have shown that time-resolved analysis of cardiac rhythmicity is a valuable tool for understanding concurrent changes in brain activity (Candia-Rivera *et al*., 2022; Sargent *et al*., 2024), highlighting the advantages of the time-resolved analysis over time-averaged measures. Animal models have provided important insights into brain-heart interactions (Hsueh *et al*., 2023; Jammal Salameh *et al*., 2024), but the exploration of cardiac changes has largely been limited to simple increases or decreases in heart rate, overlooking the contextual shifts in cardiac rhythmicity.

We recently proposed a time-resolved measurements of sympathetic and parasympathetic modulations of cardiac dynamics in humans, which was validated in different experimental conditions demonstrating its robustness (Candia-Rivera *et al*., 2025). Our method addressed a recurrent debate about the specificity of different cardiac rhythms with respect to sympathetic or parasympathetic mechanisms. The frequency domain analysis of HRV, typically relying on the estimation of low and high frequency ranges (LF, HF), has been challenged because of the inaccurate estimation of sympathetic activity within the LF range (Goldstein *et al*., 2011). To address this issue, the proposed method performs an individual-specific separation of the slow and fast HRV oscillations in combination with concurrent changes in the baseline heart rate, enabling a more precise estimation of sympathetic and parasympathetic modulations of cardiac dynamics.

This study aimed to demonstrate the effectiveness of our individual-specific approach in rats. We hypothesized that adapting the cardiac sympathetic and parasympathetic indices could accurately track short-term changes in cardiac rhythmicity during autonomic elicitation, outperforming standard spectral measures such as LF and HF. We validate the efficacy of our method using a dataset of rats exposed to dobutamine (Hazari *et al*., 2012, 2017), where dobutamine emulates a non-resting state condition similar to the physiological responses due to exercise due to sympathetic activations and parasympathetic deactivations (Sharma *et al*., 2015). We applied our method to spontaneously hypertensive rats (SHR), known for their rapid sympathetic reactivity (Shanks *et al*., 2013), and then tested on a group of Wistar-Kyoto rats (WKY). The validation of our approach in a rat model opens new avenues for advancing our understanding of cardiovascular dynamics and their relationships with ongoing neural dynamics in different clinical conditions.

## Methods

### Dataset description

Open access datasets were used in this study (Hazari *et al*., 2012, Hazari *et al*., 2017), comprising 22 rats from two different strains were included in this study: Wistar–Kyoto (WKY) and spontaneously hypertensive rats (SHR), Charles River, Wilmington, MA, USA. The rats were 12-week-old, all males, and they weighted between 300–400 g. All rats were aseptically implanted with radiotelemeters (Model TA11CTA-F40, Data Sciences International, St. Paul, MN, USA) for continuous measurement of the electrocardiogram (ECG). ECGAuto software (EMKA technologies USA, Falls Church, VA, USA) was used to visualize individual ECG signals and measure interbeat interval (IBI) or R-to-R peak segment durations (RR series).

The rats were catheterized intravenously in the left jugular vein for the administration of dobutamine. Dobutamine (Sigma-Aldrich, St. Louis, MO, USA) was infused 0.2 ml/min for 2 min in awake rats, after a 5-min baseline period (Hazari *et al*., 2012, Hazari *et al*., 2017). Administered doses were 20, 40, 80, 160 and 320 µg/kg.

Half of the cohort consisted of rats exposed to diesel exhaust, while the other half was exposed to purified air. However, our primary focus was not on comparing these two conditions. Instead, we concentrated on the instantaneous changes triggered by dobutamine administration. The effects of diesel exhaust on cardiovascular physiology remain inconclusive in the literature. Research suggests that significant effects are generally observed with long-term exposure (Bradley *et al*., 2013) or in individuals with pre-existing conditions (Carll *et al*., 2013). Diesel exhaust exposure has been linked to an increased risk of cardiac arrhythmias (Hazari *et al*., 2017; Rossi *et al*., 2021). However, these findings have not been consistently replicated in other models (Peretz *et al*., 2008; Mills *et al*., 2011).

For simplicity, the various doses of dobutamine administration were combined in the statistical analyses. Due to the substantial variability in the duration of both pre- and post-infusion periods, we established a minimum duration of 30 seconds for each. This criterion led to 41 trials for all SHR rats and 52 trials for all WKY rats.

The experimental protocols were approved by and in accordance with the guidelines of the Institutional Animal Care and Use Committee of the United States Environmental Protection Agency, Research Triangle Park, North Carolina. The data were gathered from the United States Environmental Protection Agency website (https://doi.org/10.23719/1503098). For further detail on the dataset, please refer to the original studies (Hazari *et al*., 2012, Hazari *et al*., 2017).

### Estimation of cardiac sympathetic and parasympathetic indices

The estimation of cardiac sympathetic and parasympathetic indices was based on a previously developed model for humans (Candia-Rivera *et al*., 2025). The model is based on the Poincaré plot, a method that depicts the fluctuations on the duration of consecutive IBI or RR series (Woo *et al*., 1992), as shown in Figure 1A. Notably, this method eliminates the need to resample the IBI series, unlike traditional HRV approaches, such as spectral estimations.

**Figure 1.**
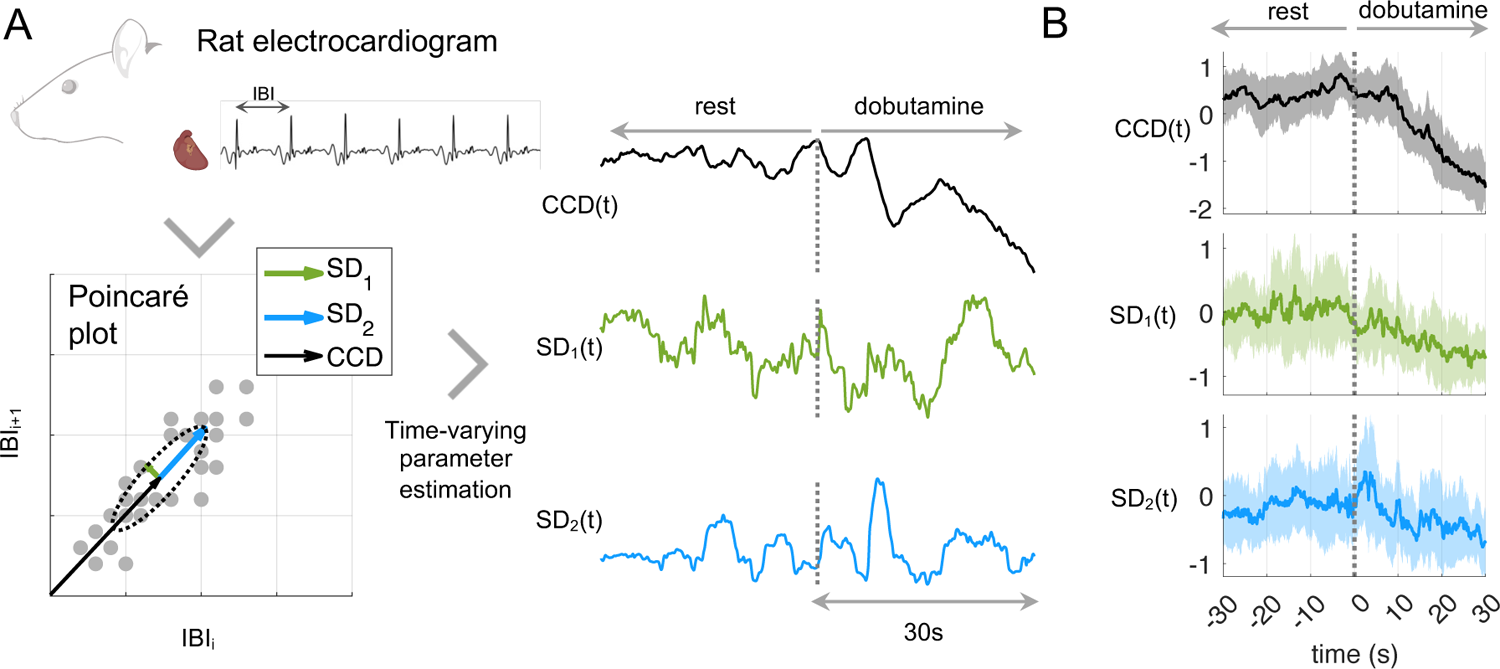
Pipeline for the estimation of the fluctuating parameters of the Poincaré plot. (A) The Poincaré plot illustrates the consecutive changes in interbeat intervals (IBI). The parameters computation includes the distance from the ellipse center to the origin (CCD), and the minor (SD1) and major ratios (SD2). (B) Group median and median absolute deviation in bold curve and shaded areas of CCD, SD1 and SD2.

Three features are extracted from the Poincaré plot: the cardiac cycle duration measured as the distance to the origin (CCD), and the minor (SD1) and major (SD2) ratios of the ellipse that quantify the short- and long-term fluctuations of HRV, respectively (Sassi *et al*., 2015).

The time-varying fluctuations of the distance to the origin and the ellipse ratios were computed with a sliding-time window, as follows:

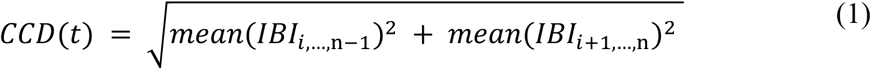

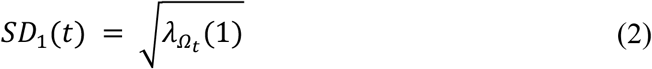

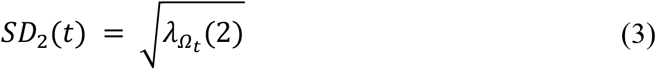

where 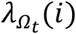 denotes the i^th^ the eigenvalue of the covariance matrix of *IBI*_i,…,n-1_ and *IBI*_i+1,…,n_, with *Ω*_*t:*_ *t* – *T* ≤ *t*_i_ ≤ *t*, and *n* is the length of IBI in the time window *Ω*_*t*_.

The original method included the implementation of four different approaches to compute the covariance matrices: “exact”, “robust”, “95%” and “approximate” (Candia-Rivera *et al*., 2025). In this study, the robust approach was used, which computes the covariance matrix using a shrinkage covariance estimator based on the Ledoit-Wolf lemma for analytic calculation of the optimal shrinkage intensity (Schäfer & Strimmer, 2005). That sliding time window to compute the indices (*T*) was fixed in 3 seconds, which takes into account that typical interbeat interval of the rat is around 100 ms and slow fluctuations occur in the range 0.1 – 1 Hz (Rowan *et al*., 2007).

The distance to the origin *CCD*_0_ and ellipse ratios *SD*_01_ and *SD*_02_ for the whole experimental duration are computed to re-center the time-resolved estimations of CCD, SD1 and SD2. The Cardiac Parasympathetic Index (*CPI*) and the Cardiac Sympathetic Index (*CSI*), are computed as follows:

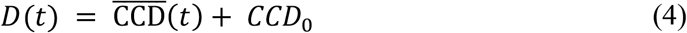

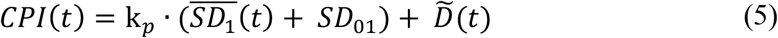

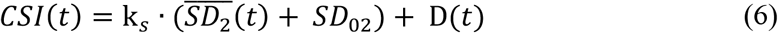

Where 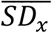 is the demeaned *SD*_*x*_ and 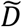 is the flipped *D* with respect the mean.

The coefficients k_*p*_ and k_*s*_ determine the influence of fast and slow HRV oscillations relative to changes in the baseline heart rate. In this study, these values were experimentally determined using the SHR dataset and subsequently validated in the WKY dataset. The coefficients were established by comparing the experimental conditions (rest vs. dobutamine) across a range of k_*p*_ and k_*s*_ values (from 1 to 20). Statistical differences were assessed based on paired comparisons using the Wilcoxon test.

### Estimation of sympathetic and parasympathetic indices using spectral components

The spectral HRV analysis was computed following the adapted Wigner–Ville method, for estimating the LF and HF time series (Orini *et al*., 2012). First, the IBI series were evenly re-sampled using spline interpolation and demeaned. The re-sampling frequency was set to 20 Hz, which is commonly set at 10 Hz or higher in rodents (Aubert *et al*., 1999). The pseudo-Wigner–Ville algorithm consists of a two-dimensional Fourier transform with an ambiguity function kernel to perform two-dimensional filtering (Orini *et al*., 2012). It is important to note that in rats, the inter beat interval (IBI) can be as short as 100 ms, compared to the typical range of 600 to 1000 ms in humans (Konopelski & Ufnal, 2016). As a result, the LF and HF bands in rat studies have been redefined multiple times, typically as 0.1-1 Hz and 1-3.5 Hz (or alternatively 0.1-1.5 Hz and 1.5-5 Hz), respectively (Rowan *et al*., 2007). In this study, the frequency bands were defined as LF: 0.1-1 Hz and HF: 1-3.5 Hz.

### Statistical analysis

To statistically evaluate the performance of the two methods (time-frequency and Poincaré plot) in discerning the experimental conditions, we used a two-sided non-parametric Wilcoxon signed-rank test for paired samples. The time-resolved information for all the estimated features was condensed as the average value for each experimental session (30 s duration each condition), and the group-wise descriptive measures are expressed as medians and median absolute deviations. Multiple comparisons were corrected using the Bonferroni rule.

Post-hoc exploratory analyses were conducted to assess the contribution of each variable—rat type (SHR or WKY), exposure (purified air or diesel exhaust), and dobutamine dose (20, 40, 80, 160 and 320 µg/kg)—as well as their potential interaction effects on the differences in CSI and CPI values. These differences were calculated as the change in values 30 seconds after dobutamine administration compared to 30 seconds before. Model performance was evaluated using the Akaike Information Criterion (AIC), assessing both the individual contributions of the variables and their interactions in fitting the model. The model with the lowest AIC value was considered the best in each case.

## Results

We investigated cardiac dynamics in two rat strains subjected to autonomic elicitation through dobutamine infusion. Our method, based on Poincaré plot descriptors of RR intervals, computed cardiac sympathetic and parasympathetic indices (CSI and CPI). Using the SHR strain, we determined parameters k_*p*_ and k_*s*_, which define the weight of HRV components with respect to the changes in baseline heart rate. Parameter definition involved testing k_*p*_ and k_*s*_, values to effectively distinguish between rest and dobutamine conditions for CSI and CPI estimation, respectively. The Z-value from the Wilcoxon test comparing both conditions served as an indicator for optimal parameters. The inflection point for CSI was identified at k_*s*_ =3, and for CPI, it was at k_*p*_=12, as illustrated in Figure 2A. Figure 2B provides an example of a single rat, depicting the estimation of CSI and CPI for various values of k_*p*_ and k_*s*_.

**Figure 2.**
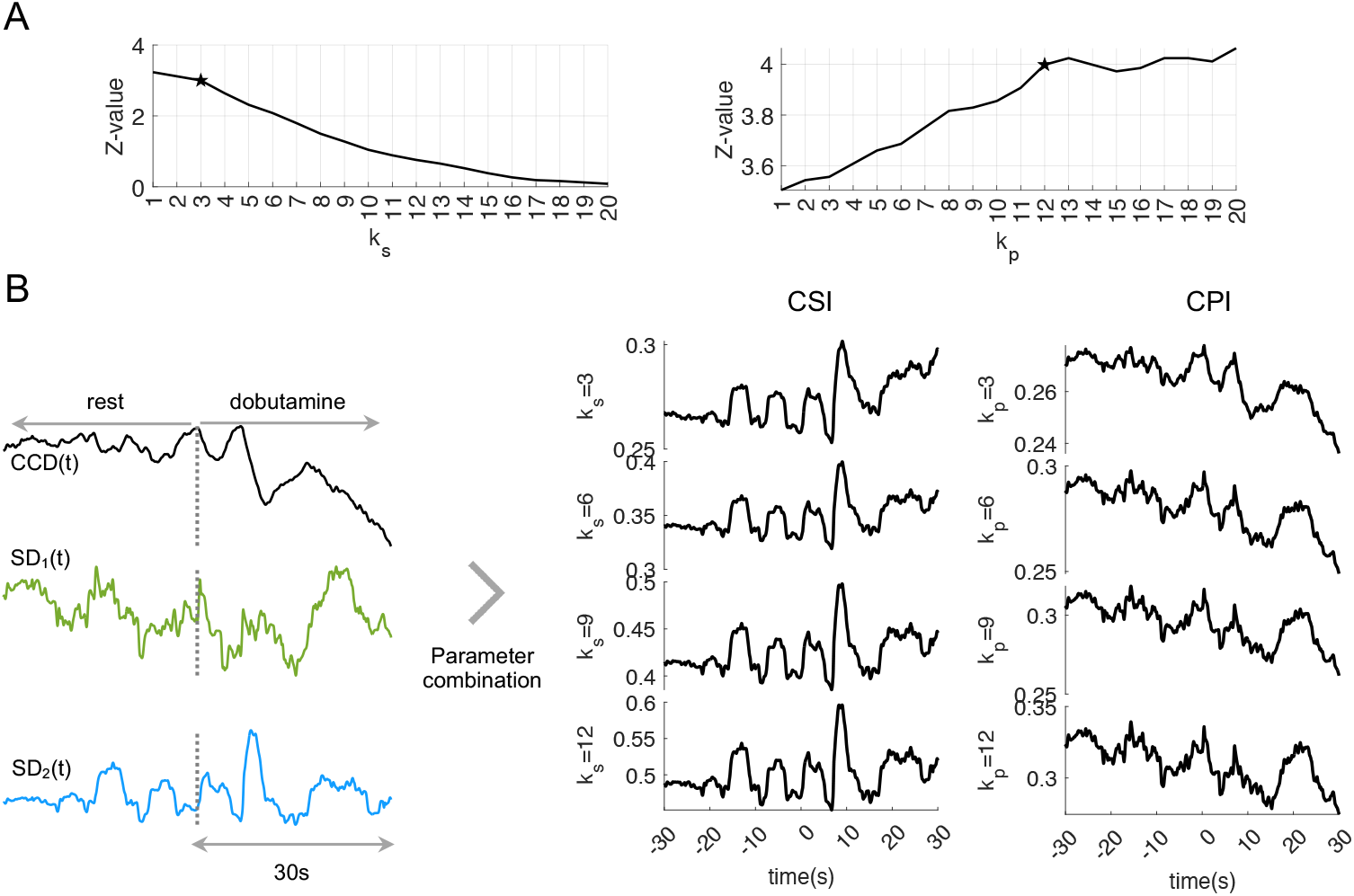
Estimation of the parameters k_*p*_ and k_*s*_ for the estimation of CSI and CPI in the SHR strain. (A) The Z-value from the Wilcoxon test comparing CSI and CPI for the conditions rest vs dobutamine, as a function of k_*p*_ and k_*s*_ values. The star indicates the inflection point from which k_*s*_ =3 and k_*p*_=12 were used for the rest of the study. (B) Changes in R, SD1 and SD2 for one rat in the transition from rest to dobutamine infusion, and their respective CSI and CPI estimation based on different values of k_*p*_ and k_*s*_.

Using the optimal parameters identified in Figure 2, we present the results of the statistical comparison between rest and dobutamine conditions for the two rat strains in Figure 3. Furthermore, we include comparisons of their spectral counterparts, LF and HF. Our findings indicate a consistently higher distinguishability of conditions for CSI and CPI compared to LF and HF.

**Figure 3.**
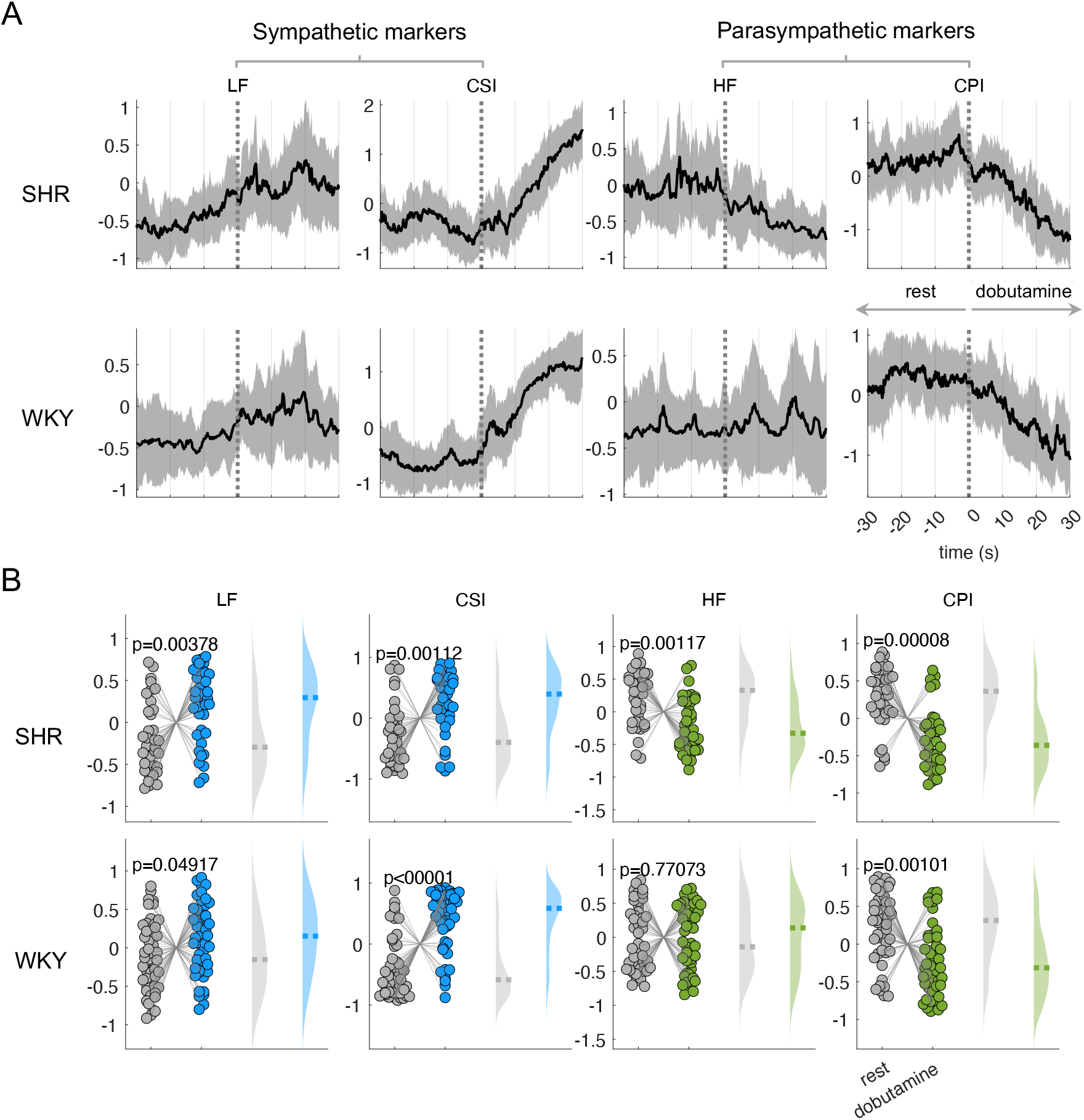
Cardiac autonomic indices CSI and CPI, and their spectral counterparts LF and HF under the rest to dobutamine transition. (A) Time course of the computed indices between −30 to 30 s with respect to the condition change onset. The plot indicates the group median and the shaded areas the median absolute deviation. Time series were z-score normalized per subject before averaging for visualization purposes. (B) Statistical comparison using a signed rank Wilcoxon test, comparing the mean 30 s after the condition change with respect to the 30 s before. Dashed lines indicate the group median. All signal amplitude units are arbitrary units.

Figure 3 also illustrates time-resolved transitions for all autonomic markers, segmented by rat strain. In Figure 3A, the SHR strain exhibits a distinct increase in sympathetic markers following dobutamine infusion, with CSI surpassing LF. Simultaneously, there is a reduction in parasympathetic activity, where CPI proves more pronounced than HF. Similar sympathetic trends are observed in the WKY strain, while the WKY strain displays no significant change in HF after dobutamine infusion, with CPI emerging as an effective marker in this scenario. The distribution of averaged autonomic markers during rest and dobutamine periods in Figure 3B further underscores the robustness of CSI and CPI in comparison to their spectral estimates’ counterparts.

As part of the exploratory analysis, various linear mixed models were fitted to examine CSI and CPI values as functions of rat type (SHR or WKY), exposure (purified air or diesel exhaust), and dobutamine dose (20, 40, 80, 160, and 320 µg/kg).

For CSI, in two-factor models (N = 1, 2, 3), the lowest AIC value was observed when the dose variable was excluded, indicating that CSI is primarily influenced by diesel exhaust exposure and rat type. In three-factor models (N = 4 to 8), the best fit was achieved when dose was treated as an independent variable alongside the interaction between diesel exhaust exposure and rat type. This suggests that dose does not significantly interact with the other variables regarding its effects on CSI.

For CPI, in two-factor models, diesel exhaust exposure and dose appeared to have a stronger influence on CPI values. In three-factor models, CPI was best explained by considering the interaction among all three factors. These findings, although limited by the number of data points, suggest that CSI and CPI are influenced differently by the analyzed factors. Further studies with larger datasets are needed to confirm these observations.

**Table 1.** Linear Mixed Model fitting Cardiac Sympathetic Index (CSI)

| N model | Formula | AIC |
| --- | --- | --- |
| **1** | **CSI ~ 1 + Exposure + RatType + (1\|ID)** | **-406.7596** |
| 2 | CSI ~ 1 + Exposure + Dose + (1\|ID) | -368.4319 |
| 3 | CSI ~ 1 + RatType + Dose + (1\|ID) | -366.8458 |
| 4 | CSI ~ 1 + Exposure + RatType + Dose + (1\|ID) | -376.7366 |
| **5** | **CSI ~ 1 + Dose + Exposure\*RatType + (1\|ID)** | **-498.3792** |
| 6 | CSI ~ 1 + RatType + Exposure*Dose + (1\|ID) | -346.9463 |
| 7 | CSI ~ 1 + Exposure + RatType*Dose + (1\|ID) | -341.2821 |
| 8 | CSI ~ 1 + Exposure * RatType * Dose + (1\|ID) | -403.9011 |

ID={1, ..., 22}, Exposure={Air, Diesel}, RatType={SHR, WKY}, Dose={20, 40, 80, 160, 320 μg/kg}. AIC: Akaike Information Criterion (lowest value in bold indicates best fitting)

**Table 2.** Linear Mixed Model fitting Cardiac Parasympathetic Index (CPI)

| N model | Formula | AIC |
| --- | --- | --- |
| 1 | CPI ~ 1 + Exposure + RatType + (1\|ID) | 85.0815 |
| **2** | **CPI ~ 1 + Exposure + Dose + (1\|ID)** | **84.1944** |
| 3 | CPI ~ 1 + RatType + Dose + (1\|ID) | 85.9075 |
| 4 | CPI ~ 1 + Exposure + RatType + Dose + (1\|ID) | 78.1941 |
| 5 | CPI ~ 1 + Dose + Exposure*RatType + (1\|ID) | 62.9700 |
| 6 | CPI ~ 1 + RatType + Exposure*Dose + (1\|ID) | 59.6205 |
| 7 | CPI ~ 1 + Exposure + RatType*Dose + (1\|ID) | 63.8296 |
| **8** | **CPI ~ 1 + Exposure \* RatType \* Dose + (1\|ID)** | **-44.6750** |

ID={1, ..., 22}, Exposure={Air, Diesel}, RatType={SHR, WKY}, Dose={20, 40, 80, 160, 320} μg/kg. AIC: Akaike Information Criterion (lowest value in bold indicates best fitting)

## Discussion

Heart rhythmicity analysis has the potential to be a valuable noninvasive technique for assessing autonomic nervous activity in rats (Kuwahara *et al*., 1994). Standard time-averaged HRV measures often fail to capture ongoing physiological dynamics because the averaging process, typically spanning a few minutes, masks fluctuations that occur at the level of seconds. For instance, invasive recordings on the left cervical vagus nerve in rats have revealed that tonic vagal activity does not correlate with commonly used HRV metrics (Marmerstein *et al*., 2021). This indicates the need for further development of methods to understand the modulation of sympathetic and parasympathetic activity on cardiac dynamics.

We developed and validated a method using a rat heart dataset, where pharmacological autonomic elicitation was induced with dobutamine. This drug, a positive inotropic agent, is known to mimic sympathetic activation and parasympathetic deactivation, as observed in conditions like physical exercise (Mahoney *et al*., 2016). In our proposed method, previously validated in humans (Candia-Rivera & Chavez, 2024*a*, 2024*b*; Candia-Rivera *et al*., 2025), we accurately show the time-dependent effects of autonomic elicitation induced by dobutamine in rats, unraveling some of the key components of cardiac regulation by both sympathetic and parasympathetic systems. As expected, our results show that dobutamine triggers an increase in sympathetic activity, shown as a sudden increase in slow fluctuations in HRV, followed by a transient increase in the baseline heart rate. In parallel, a transient decrease in parasympathetic activity occurs. We did not observe notable differences on the effects in the WKY with respect to the SHR strain, although sympathetic hyperreactivity is expected in the SHR strain, as compared to WKY (Luft *et al*., 1986). However, the sympathetic reactivity has been reported only in changes in the baseline heart rate, and not necessarily in the slow fluctuations in HRV (Ricksten *et al*., 1984). Importantly, our approach surpasses the spectral counterparts, LF and HF, in the description of sympathetic and parasympathetic activities, respectively.

Our method offers various advantages, as compared to state-of-the-art methods. First, it is based on the Poincaré plot, which is a computationally efficient tool to depict the beat-to-beat alterations in IBIs (Woo *et al*., 1992; Brennan *et al*., 2001). Second, it considers parallel autonomic modulations in both baseline heart rate and HRV. It is important to study baseline heart rate in conjunction with HRV to fully comprehend sympathetic dynamics (Sacha *et al*., 2013), as sympathetic modulations of heartbeat dynamics may be reflected solely in changes in the baseline heart rate, rather than changes in HRV (e.g., LF or LF/HF ratio), and vice versa (Gonçalves *et al*., 2010). Third, our proposal focuses on developing a time-resolved estimation method, which could reduce the need for several minutes of averaging to assess a single experimental condition, but also enabling a comprehensive exploration of the dynamic shifts in autonomic regulations occurring in parallel with other organs, such as the brain (Samuels, 2007; Bashan *et al*., 2012; Silvani *et al*., 2016), This underscores the importance of modeling brain-heart interactions to better understand multisystem dysfunctions (Chen *et al*., 2021).

Further exploration is needed to elucidate the dynamics of rat cardiac activity that extend beyond baseline heart rate or variations in HRV. Specially in the context of diesel exhaust exposure, which can potentially trigger cardiac arrythmias that can be capture as nonlinear patterns of activity (Hazari *et al*., 2017; Rossi *et al*., 2021). Linear mixed models suggest that cardiac sympathetic activity is influenced by the interaction between diesel exhaust exposure and the SHR rat type. While the available data may be limited, we cannot exclude the possibility that diesel exhaust exposure induces short-term changes in cardiac sympathetic or parasympathetic indices, potentially driven by cardiac conduction instability and inflammatory processes (Hazari *et al*., 2017; Rossi *et al*., 2021). Nonlinear methods have been suggested as an alternative for investigating autonomic dynamics. However, these methods have produced conflicting results, suggesting that increased sympathetic modulation may contribute to nonlinear dynamics (Silva *et al*., 2017), while also indicating that increased sympathetic modulation is likely the main cause of lower entropy (Silva *et al*., 2016).

Although our study has some limitations, such as a small sample size or the inclusion of all-male strains, the results demonstrate the effectiveness of our estimation of autonomic system parameters. Another limitation of this study is the inability to determine whether specific dobutamine doses or their cumulative effects after repeated administrations influence CSI or CPI values. Linear mixed models revealed that CSI was best explained by the interaction between exposure and rat type, while CPI was best explained by the interaction among all three factors studied. These findings suggest that a dose effect, although minimal, cannot be ruled out. Such an effect would likely manifest as rapid sympathetic and parasympathetic modulation of the heart, occurring immediately after a perturbing element, within a complex interplay of regulatory loops spanning multiple timescales (Pathak *et al*., 2006).

Our proposal has not been tested under other standard conditions of autonomic elicitation, such as monitoring heart rate during cold exposure (Kuwahara *et al*., 2009), which constitutes the main limitation of our study. However, our proposal constitutes a first attempt for a developing field of research on time-resolved brain-heart analysis, applied to animal models. It is worth mentioning that the study under pharmacological interventions to modulate autonomic nervous system activity remain a standard approach (Coleman, 1980; Kuwahara *et al*., 1994; Mangin *et al*., 1998; Aubert *et al*., 1999; Ohnuki *et al*., 2001; Ramaekers *et al*., 2002*a*; Beckers *et al*., 2006; Bernat *et al*., 2023).

For accurate time-resolved estimation, it is crucial to set a sliding time window that captures cardiac rhythm oscillations. We established a 3-second window to balance high time resolution with the ability to capture slow fluctuations in HRV, a key marker of sympathetic activity. In future studies, determining this window size should rely on the known physiological priors, considering expected response latency and the involved frequencies.

Our method opens new research opportunities for clinical and translational studies to explore the potential of using heartbeat dynamics as biomarkers for assessing conditions characterized by disrupted autonomic dynamics. In severe conditions such as cardiogenic shock or heart failure, a significant alteration in arterial tension can critically compromise hemodynamics, potentially leading to organ failure. To optimize organ perfusion, sympathomimetic agents like dobutamine, a β1-adrenergic agonist, have been developed to enhance heart contractility and output (Dubin *et al*., 2017). However, a comprehensive understanding of this intricacy requires further rigorous investigations in well-controlled in vivo settings.

Future research may include the description of our developed markers under different stressors (Ramaekers *et al*., 2002*b*; Inagaki *et al*., 2004; Raffai *et al*., 2006; Park *et al*., 2017), and cardiovascular conditions (Henze *et al*., 2008; Scridon *et al*., 2012; Han *et al*., 2014). Further research could deepen our understanding of the fluctuations of cardiac rhythmicity in the context of circadian rhythms, which are typically limited to describing long-term changes in the baseline heart rate alone (Van Den Buuse, 1994).

Our development may also enable the investigation of disruptions that may impact the connection between the brain and heart, and their subsequent effects on neural functions. For instance, in the study of severe brain damage resulting from cardiovascular conditions like stroke (Kodama *et al*., 2018) or anoxia (Gonçalves *et al*., 2008), as well as in cases of damage to the spinal cord (Inskip *et al*., 2012). These conditions present compelling areas of interest for further exploration in which the time-resolved estimation of autonomic dynamics can be a hallmark in the comprehension of nervous-system-wise complex dynamics.

## Conclusions

Our study presents a robust approach for time-resolved estimation of sympathetic and parasympathetic modulations in rat cardiac dynamics, demonstrating its efficacy in capturing autonomic responses triggered by pharmacological interventions. The application of our method in animal models paves the way for future research aimed at unraveling the complex interplay between cardiac and brain activity in various pathological conditions.

## Acknowledgements

This research was supported by the European Commission, Horizon MSCA Program (grant n° 101151118).

## Author contributions

Conceptualization: DCR; Figures: DCR; Data Analysis: DCR; Writing – first version of the manuscript: DCR; Writing – review & editing: DCR, SCF, FDVF, MC.

